# Physiological Roles of an *Acinetobacter*-specific σ Factor

**DOI:** 10.1101/2024.07.08.602572

**Authors:** Emily E. Bacon, Kevin S. Myers, Rubén Iruegas-López, Amy B. Banta, Michael Place, Ingo Ebersberger, Jason M. Peters

## Abstract

The Gram-negative pathogen *Acinetobacter baumannii* is considered an “urgent threat” to human health due to its propensity to become antibiotic resistant. Understanding the distinct regulatory paradigms used by *A. baumannii* to mitigate cellular stresses may uncover new therapeutic targets. Many γ-proteobacteria use the extracytoplasmic function (ECF) σ factor, RpoE, to invoke envelope homeostasis networks in response to stress. *Acinetobacter* species contain the poorly characterized ECF “SigAb;” however, it is unclear if SigAb has the same physiological role as RpoE. Here, we show that SigAb is a metal stress-responsive ECF that appears unique to *Acinetobacter* species and distinct from RpoE. We combine promoter mutagenesis, motif scanning, and ChIP-seq to define the direct SigAb regulon, which consists of *sigAb* itself, the stringent response mediator, *relA*, and the uncharacterized small RNA, “*sabS*.” However, RNA-seq of strains overexpressing SigAb revealed a large, indirect regulon containing hundreds of genes. Metal resistance genes are key elements of the indirect regulon, as CRISPRi knockdown of *sigAb* or *sabS* resulted in increased copper sensitivity and excess copper induced SigAb-dependent transcription. Further, we found that two uncharacterized genes in the *sigAb* operon, “*aabA*” and “*aabB*”, have anti-SigAb activity. Finally, employing a targeted Tn-seq approach that uses CRISPR-associated transposons, we show that *sigAb*, *aabA*, and *aabB* are important for fitness even during optimal growth conditions. Our work reveals new physiological roles for SigAb and SabS, provides a novel approach for assessing gene fitness, and highlights the distinct regulatory architecture of *A. baumannii*.

**Importance:** *Acinetobacter baumannii* is a hospital-acquired pathogen, and many strains are resistant to multiple antibiotics. Understanding how *A. baumannii* senses and responds to stress may uncover novel routes to treat infections. Here, we examine how the *Acinetobacter*-specific transcription factor, SigAb, mitigates stress. We find that SigAb directly regulates only a small number of genes, but indirectly controls hundreds of genes that have substantial impacts on cell physiology. We show that SigAb is required for maximal growth, even during optimal conditions, and is acutely required during growth in the presence of elevated copper. Given that copper toxicity plays roles in pathogenesis and on copper-containing surfaces in hospitals, we speculate that SigAb function may be important in clinically-relevant contexts.

## Introduction

The Gram-negative γ-proteobacterium, *Acinetobacter baumannii*, is a nosocomial pathogen with the ability to cause severe infections such as pneumonia and bacteremia (1). Its classification as an “urgent threat” to human health by the Centers for Disease Control stems from isolates that are resistant to nearly all clinically relevant antibiotics (2). At least some of this resistance can be attributed to the high abundance and activity of efflux pumps encoded in the *A. baumannii* genome (3, 4) among other resistance elements (5). Transcriptional regulation of these pumps is complex and incompletely understood (6).

Sigma (σ) factors are critical components of bacterial transcription regulation that direct RNA polymerase (RNAP) to specific promoters (7). Extracytoplasmic function (ECF) σ factors are a type of alternative σ factor that can play roles in cell homeostasis during unstressed growth but can also be activated under specific environmental conditions including envelope, oxidative, and metal stresses, among others (8–11). ECFs control downstream stress responses by directing RNAP to specialized subsets of promoters (regulon); thus, activating genes involved in mitigating the stress. Structurally, ECF σ factors contain two globular domains for binding promoter elements—σ_2_ and σ_4_ interact with the −10 and −35 promoter elements, respectively—and require near-consensus promoters due to their reduced capacity for promoter melting (12, 13).

RpoE is one of the most extensively studied ECFs (14, 15). In *Eschericha coli*, the RpoE signal transduction pathway is activated upon detecting envelope stress in the form of misfolded outer membrane proteins or lipopolysaccharide (LPS) intermediates. These molecules are recognized by periplasmic proteases or the anti-σ RseB which accelerates proteolysis of the anti-σ RseA, releasing RpoE to transcribe its regulon (16, 17). The RpoE regulon includes over 100 protein coding genes as well as 3 non-coding RNAs that are critical to its envelope homeostasis function (18–20). Many γ-proteobacteria, such as *Pseudomonas aeruginosa* and *Vibrio* species, contain a homolog of RpoE, which recognize similar promoter sequences and regulate overlapping sets of genes (e.g., LPS transport and outer membrane repair genes) (21–24). *A. baumannii* strains contain ECF sigma factors (25), but their functions are poorly characterized. Despite the importance of RpoE to envelope homeostasis in γ-proteobacteria, it is unclear if any ECF σ factors in *A. baumannii* play similar roles.

Gene regulatory patterns and regulators in *A. baumannii* are often distinct from those observed in model organisms, such as *E. coli* K-12. For example, *A. baumannii* lacks homologs of key stress response genes including the general stress response σ factor RpoS and the Rcs envelope stress signal transduction system (26, 27). Instead, *A. baumannii* encodes the two-component system BfmRS that exhibits phenotypic overlap with Rcs and other envelope stress responses (27, 28). Additionally, *A. baumannii* contains numerous genes of unknown function, including putative transcription factors and cell envelope genes (25, 29). *A. baumannii* can survive a wide range of stresses including prolonged desiccation on surfaces, metal toxicity and oxidative stress during host infection, and evading antibiotic killing (30–32). However, little is known if *A. baumannii* ECF σ factors play a role in mitigating these stresses.

Here, we investigate the regulon and physiological roles of an *Acinetobacter*-specific ECF σ factor we call, “SigAb”. The gene encoding SigAb (ACX60_04565 in ATCC 17978) has been annotated as “RNA polymerase sigma factor”, or “*sigX*”, and was suggested to be similar to RpoE and the *Pseudomonas* homolog, AlgU, based on the small number of residues that align in predicted σ_2_ and σ_4_ domains (25). We find that SigAb recognizes a distinct DNA binding site and regulon from RpoE, and we show that SigAb instead has roles in metal resistance and general fitness during growth without added stressors. Finally, we discuss the implications of our work for gene regulation in *Acinetobacter* species.

## Results

### SigAb is an *Acinetobacter*-specific σ factor

We first sought to compare SigAb to characterized ECF σs, including RpoE. As expected, a search for proteins with similar predicted folds using Phyre2 (33) returned high-confidence matches to structurally characterized ECFs (Fig. S1). Further, we were able to model SigAb in place of *E. coli* RpoE in an RpoE-RNAP holoenzyme structure (Fig. S2 (34)). However, SigAb showed low overall primary sequence identity to *E. coli* RpoE (18%), the RpoE ortholog in *P. aeruginosa* AlgU (19%), and the *P. aeruginosa* ECF SigX (20%), and key DNA binding residues in RpoE differed in SigAb (e.g., RpoE F64, R76, S172, F175, etc.), suggesting distinct interactions with promoter DNA.

The low sequence identity between RpoE and SigAb left their precise evolutionary relationships unclear. To further shed light on the evolutionary trajectory that gave rise to these sequences, we determined the phylogenetic profiles of SigAb and RpoE using a targeted ortholog search. Interestingly, this suggested at first sight that RpoE and SigAb are indeed orthologs as their phylogenetic profiles contain, in part, the same proteins. To test this hypothesis further, we created a non-redundant protein list from the two profiles and selected a representative set covereing the γ-proteobacterial orders. A subsequent multiple sequence alignment revealed a conspicuous conservation pattern (Fig. 1a). Sequences from a diverse set of orders including the Enterobacterales, Vibrionales, Pseudomonadaceae, Pasteurellales, and Alteromonadales are highly conserved; Among these sequences, we find RpoE of *E. coli*.

**Figure 1.**
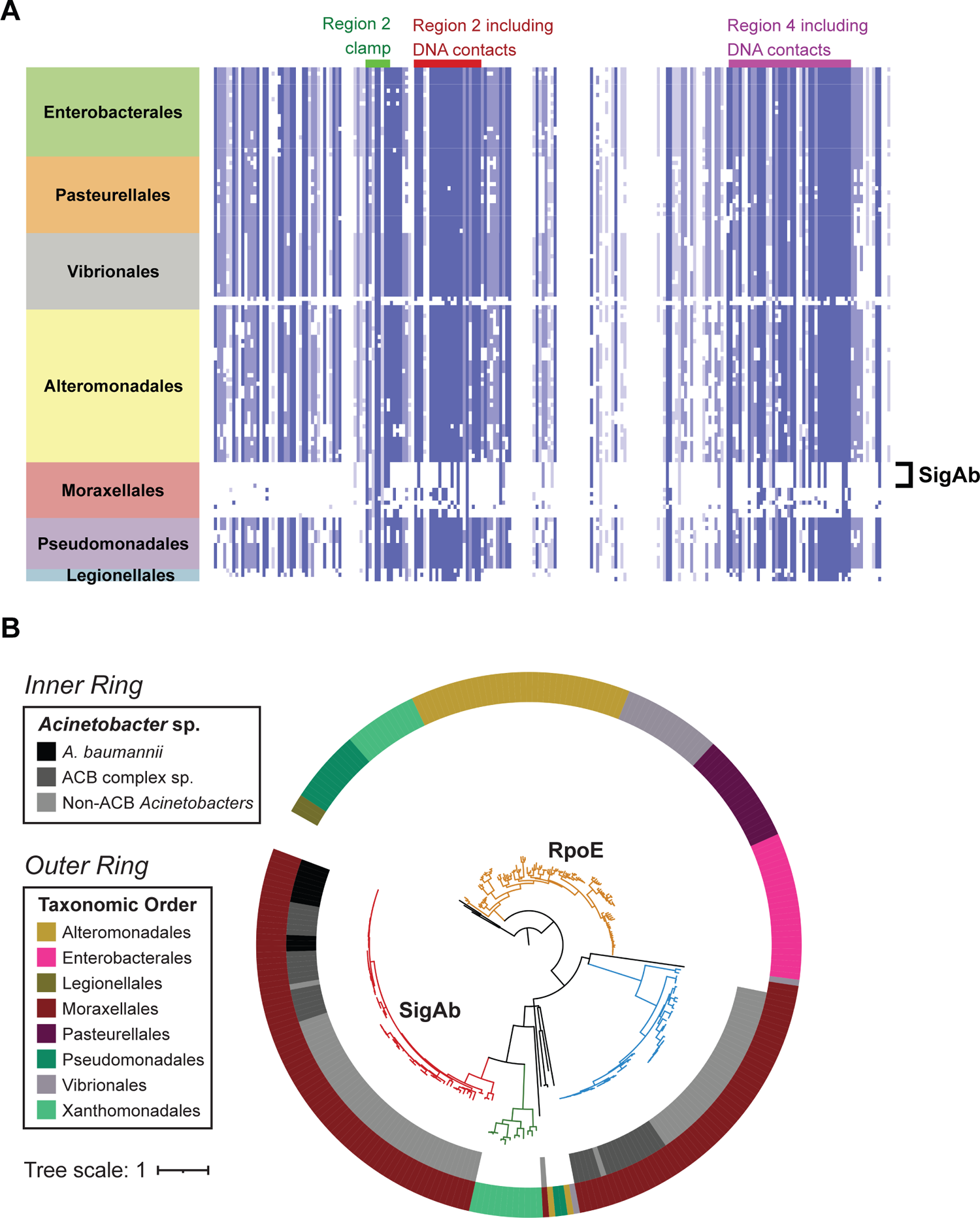
SigAb is an *Acinetobacter*-specific σ factor. **A** Alignment of ECF σ factors from SigAb ortholog search across the γ-proteobacteria. Alignment is colored by protein sequence identity, with darker blue indicating greater amino acid conservation. The Moraxellales family contains *Acinetobacter* species. **B** Phylogenetic tree based on alignment from (a). SigAb orthologs are in red and RpoE orthologs are in gold. An additional ECF distinct from SigAb and RpoE present in non-*baumannii Acinetobacters* is in blue. Outer ring denotes γ-proteobacterial taxonomic order, and inner ring distinguishes between different *Acinetobacter* sp. classifications, including the *A. calcoaceticus-baumannii* (ACB) complex.

Moraxellaceae—and in particular members of the genus *Acinetobacter*—formed a separate group of sequences including SigAb whose conservation pattern is clearly distinct from RpoE (Fig. 1a). We next constructed a phylogenetic tree based on our alignment (Fig. 1b). We found that the conservation pattern seen in the multiple sequence alignment is reflected in the tree topology. Sequences from different γ-proteobacterial orders are grouped into one clade to the exclusion of the representatives from the genus *Acinetobacter.* This placement is at odds with the accepted evolutionary relationships of the organisms where the Moraxellales are considered the next relatives of the Pseudomonadales. It strongly suggests that SigAb in *Acinetobacter* represents a distinct evolutionary lineage from that of RpoE. Furthermore, RpoE-like ECFs are absent from the *Acinetobacter* genomes we queried, including *A. baumannii* and other members of the *A. calcoaceticus-baumannii* (ACB) complex. Taken together, SigAb is an ECF σ factor found in *Acinetobacter* species that is evolutionarily distinct from RpoE.

### SigAb-dependent promoters are distinct from those recognized by RpoE

Protein modeling and evolutionary analysis highlighted distinctions between RpoE and SigAb, raising the possibility that SigAb could recognize a different promoter sequence.

Because ECF σ factor expression is often autoregulated, we investigated the DNA sequence upstream of *sigAb* for a possible SigAb promoter. Indeed, we found an upstream sequence that was specifically recognized by SigAb (P*_sigAb_*) (Fig. 2). To map P*_sigAb_*, we used 5′ RACE to determine the 5′ end of the *sigAb* transcript (Fig. S3a). We next aligned the DNA sequence upstream of the putative P*_sigAb_* transcription start site (TSS) across selected *Acinetobacter* species, finding highly conserved motifs that could potentially serve as promoter −10, −35, and UP elements (35) (Fig. S3b). To test for SigAb-dependent promoter activity, we cloned the putative P*_sigAb_* sequence upstream of a Red Fluorescent Protein reporter gene (*mRFP*) and integrated the reporter into the genomes of *A. baumannii*, which contains a native copy of *sigAb*, and *E. coli*, which lacks *sigAb* (Fig. 2a). We found that overexpression (OE) of SigAb from a multi-copy plasmid increased reporter activity by >150-fold in *A. baumannii* (Fig. 2b) and that the presence of the *sigAb* gene was necessary and sufficient for P*_sigAb_* reporter activity in *E. coli* (Fig. 2c). The absence of reporter activity in *E. coli* lacking *sigAb* suggests that RpoE does not recognize P*_sigAb_* (Fig. 2c). Further, a P*_rpoE_* reporter showed activity in *E. coli*, but not *A. baumannii* (Fig. S4), supporting that SigAb does not recognize RpoE-dependent promoters.

**Figure 2.**
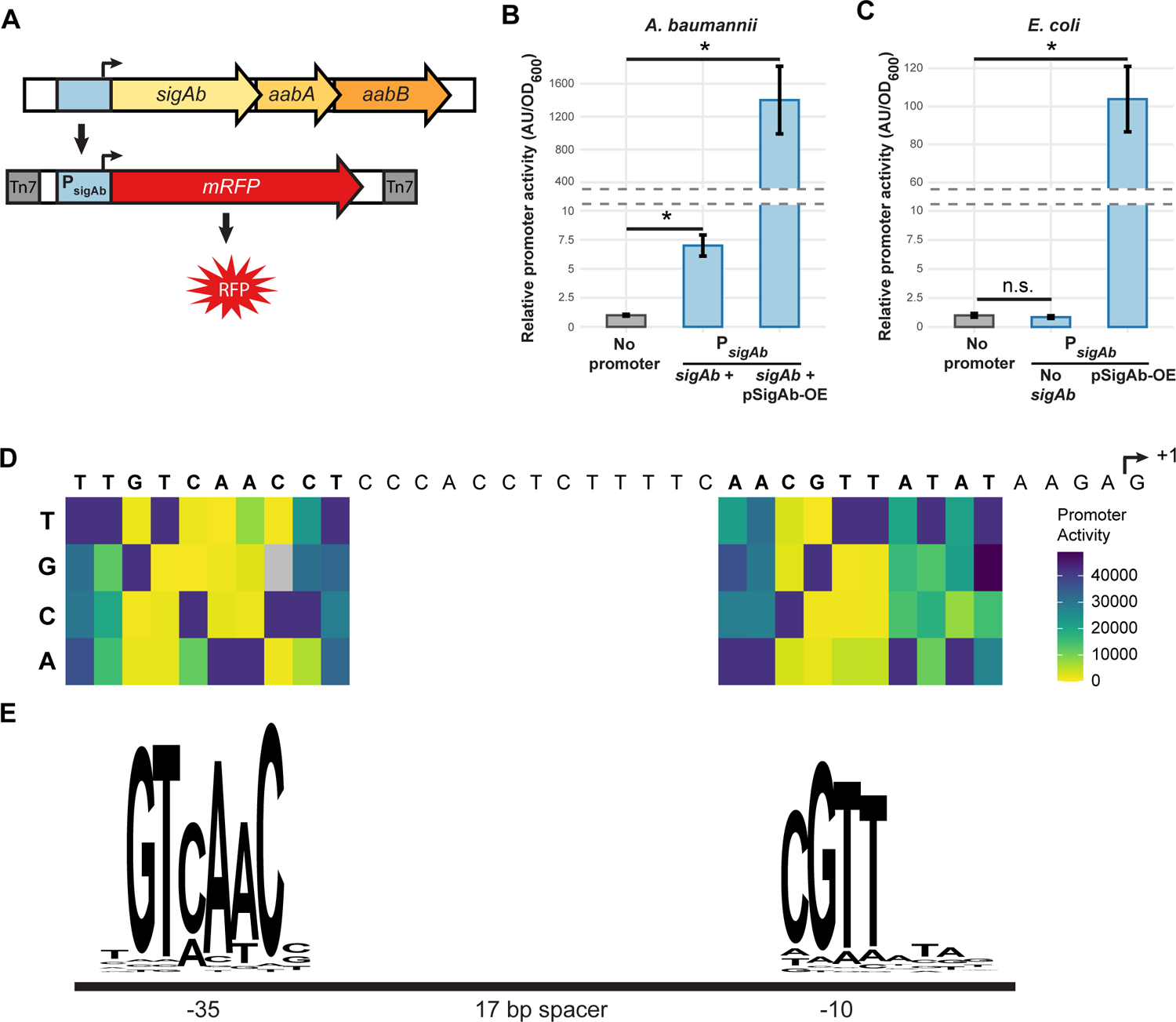
Identification of core promoter sequence recognized by SigAb. **A** mRFP fluorescent reporter to assay *sigAb* promoter (P*_sigAb_*) activity. Reporter is stably integrated in the chromosome in the *att*_Tn*7*_ site. *aabA* and *aabB* encoded downstream of *sigAb* are predicted to be in an operon. **B and C** P*_sigAb_*-*mrfp* reporter activity in WT or *sigAb* overexpression strains in (b) *A. baumannii* or (c) *E. coli*. Promoter activity is calculated as absorbance units (AU) normalized to OD_600_ and the no promoter control (n=3). Data are represented as the mean ± s.d. and significance was calculated with a two-tailed Student’s *t*-test (p<0.05). **D** Systematic mutagenesis of P*_sigAb_* sequence. TSS was identified using 5’ RACE. Point mutations in promoter were assayed for activity using mRFP reporter. Heatmap shown is median of n=5 assays for 1-13 biological replicates per mutation. **E** Quantification of P*_sigAb_* mutagenesis data from (d) as an activity logo.

With a validated reporter in hand, we sought to determine which bases within P*_sigAb_* are important for SigAb activity. Conserved positions in our P*_sigAb_* alignment across *Acinetobacter* species provided a starting point for systematic mutagenesis of the promoter sequence. Using our P*_sigAb_* reporter, we comprehensively mutated individual bases in the putative −10 and −35 elements and measured reporter expression in *A. baumannii*; this allowed us to identify the key bases for promoter activity (Fig. 2d and S5a). We weighted P*_sigAb_* variants by promoter activity and created a SigAb activity logo, which revealed distinct −10 and −35 elements (Fig. 2e). The core SigAb −10 (CGTT) and −35 (GTCAAC) identified by our mutagenesis approach differ from those determined by promoter alignments for RpoE (−10 ∼TCAAA and −35 ∼GGAACTT (19)).

Other promoter features also contributed to P*_sigAb_* activity. P*_sigAb_* has a 17-bp spacer sequence between the −10 and −35; reducing the spacer length to 16 had little impact on activity but increasing the spacer length to 18 reduced activity by ∼3-fold (Fig. S5b). Our P*_sigAb_*alignment also suggested that a conserved run of four T bases in the spacer could impact activity.

Consistent with this, we found a modest 2.5-fold reduction in activity when all four T bases were substituted with G bases (Fig. S5c). Finally, a run of A/T bases upstream of the −35 may serve as an UP element, as substitution of this sequence with random bases reduced P*_sigAb_*activity by 10-fold (Fig. S5c). In sum, we identified a SigAb-dependent promoter, systematically defined a promoter activity motif with key sequences required for activity, and showed that this promoter is distinct from that of RpoE.

### SigAb directly controls a small regulon

As the SigAb regulon remained uncharacterized, we set out to identify direct targets of SigAb, taking a two-pronged approach: 1) we scanned the *A. baumannii* ATCC 17978 genome for putative SigAb binding sites that matched our promoter activity motif, and 2) we performed chromatin immunoprecipitation followed by sequencing (ChIP-seq) to find DNA sites occupied by SigAb in whole cells. We found that SigAb directly controls a small regulon of genes that includes one or more non-coding RNAs. We first scanned the ATCC 17978 genome for exact matches to the P*_sigAb_* −10 (CGTT) and −35 (GTCAAC) elements with spacer lengths between 16 and 18 bases. We identified a total of 17 motifs (Fig. S6a); surprisingly only three were upstream of annotated protein coding genes in an orientation that would be expected to drive downstream transcription. We individually cloned the 17 motifs into our mRFP reporter construct, finding that many of the motifs had substantial activity upon SigAb OE from a multi-copy plasmid (Fig. S6b). In addition to P*_sigAb_*, SigAb-dependent promoters were identified upstream of the gene encoding the (p)ppGpp synthetase, *relA*, a global regulator during nutrient limitation (36–38) as well as the sulfate transporter operon, *cysTW*. We found another highly active SigAb-promoter upstream of a putative small RNA (sRNA) that had previously been identified by sequencing of RNAs (RNA-seq), but the promoter for this sRNA had not been characterized (39). We call this sRNA “*sabS*” for SigAb-dependent sRNA. Importantly, the *sigAb*, *relA*, and *sabS* promoter motifs exhibit significantly greater SigAb-dependent activity than the other putative motifs (Fig. S6b).

We next used ChIP-seq to identify SigAb binding sites in whole cells grown in rich medium. For this purpose, we generated an N-terminally Halo-tagged variant of SigAb that we confirmed retained activity using our P*_sigAb_*reporter (Fig. S7). Halo-tagged proteins covalently bind Halolink resin, enabling stringent washing conditions that remove non-specific DNA (40). Peak calling analysis of three independent ChIP-seq samples showed only three sites that were both significantly enriched across at least two replicates and also contained a putative *sigAb* promoter motif (Fig. 3a, Table S5). These three enriched sites were in front of the following genes: 1) *sigAb* (Fig. 3b), 2) *relA* (Fig. 3c), and 3) *sabS* (Fig. 3d). Although *sigAb* and *sabS* showed much stronger ChIP-seq signal than *relA*, all three motifs exhibited similar induction upon plasmid-based SigAb OE in our mRFP reporter assay (Fig. 3e and Fig. 2b). We conclude that SigAb directly controls a small regulon including itself, *relA*, and one or more non-coding RNAs.

**Figure 3.**
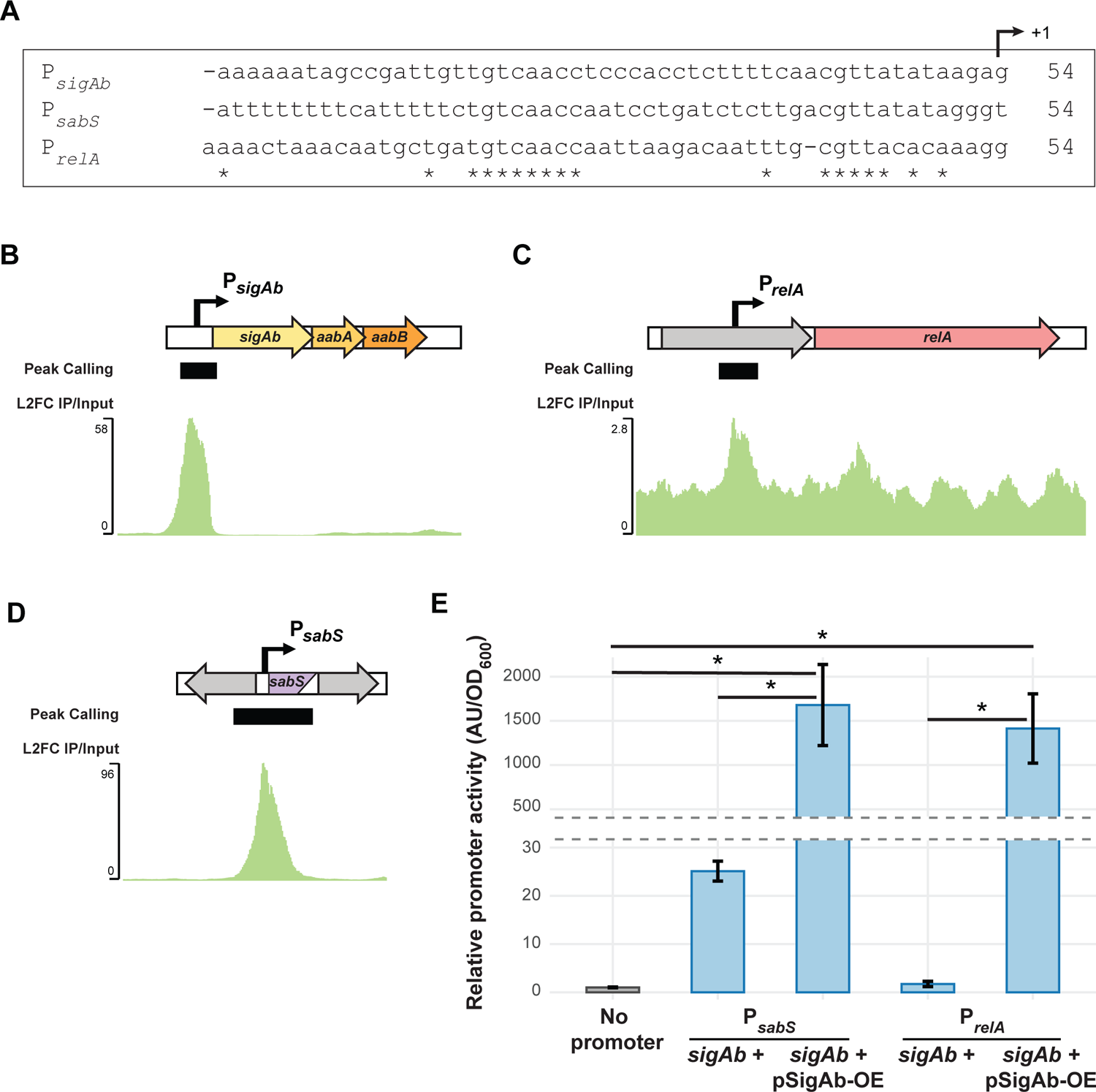
SigAb directly controls a small regulon. **A** Sequence alignment of the SigAb-dependent promoter motifs for *sigAb*, *sabS*, and *relA*. Stars indicate conserved bases and +1 indicates the putative transcription start site (TSS). **B** ChIP-seq peak at *sigAb* locus from an *A. baumannii* strain harboring HaloTagged SigAb. Data are represented as log_2_(fold change) of immunoprecipitated sample normalized to input control. Peak calling algorithm was used for significant peak identification. **C** ChIP-seq peak at the *relA* locus. **D** ChIP-seq peak at an intergenic region containing an uncharacterized sRNA, *sabS*. **E** P*_sabS_*-*mrfp* and P*_relA_*-*mrfp* reporter activity in WT or *sigAb* overexpression strains in *A. baumannii*. Relative promoter activity is calculated as absorbance units (AU) normalized to OD_600_ and the no promoter control (n=3). Data are represented as the mean ± s.d. and significance was calculated with a two-tailed Student’s *t*-test (p<0.05).

### SigAb indirectly affects global transcription

SigAb control of the global regulator *relA* and putative sRNAs raised the possibility that increases in SigAb activity could affect transcription beyond its small, direct regulon. To test for a global effect of SigAb on transcription, we overexpressed *sigAb* from the IPTG-inducible *trc* promoter on a multi-copy plasmid and performed an RNA-seq timecourse post induction (Fig. 4a). Interestingly, hundreds of genes increased in expression by 1 hour post *sigAb* induction compared to a vector only control (Fig. 4b). After only 5 minutes of induction, 125 genes had increased expression by over 2-fold (FDR < 5%), suggesting that upregulation of these genes is a secondary effect of SigAb OE (Fig. S8a). Gene set enrichment analysis showed that a variety of cellular pathways were upregulated at the 5 min timepoint including “metabolism and oxidoreductase activity,” which contained *relA* among other metabolic genes and “membrane and regulation of cellular processes” in addition to TetR and LysR-type transcription factors which may partially explain the large number of genes affected by 60 min (Fig. 4c); however, these upregulated genes lacked known RpoE targets. Although not a defined enrichment group, we note that many prophage genes (20 genes) were upregulated by SigAb OE (identified using Phaster (41)), which may be due to stress-induced prophage expression. The vast majority of these upregulated genes had no associated SigAb binding site, suggesting a widespread indirect effect of SigAb OE. Two enriched groups of SigAb upregulated genes contained transporter-encoding genes with possible relevance to *A. baumannii’*s resistance and pathogenesis lifestyles (Fig. 4d).

**Figure 4.**
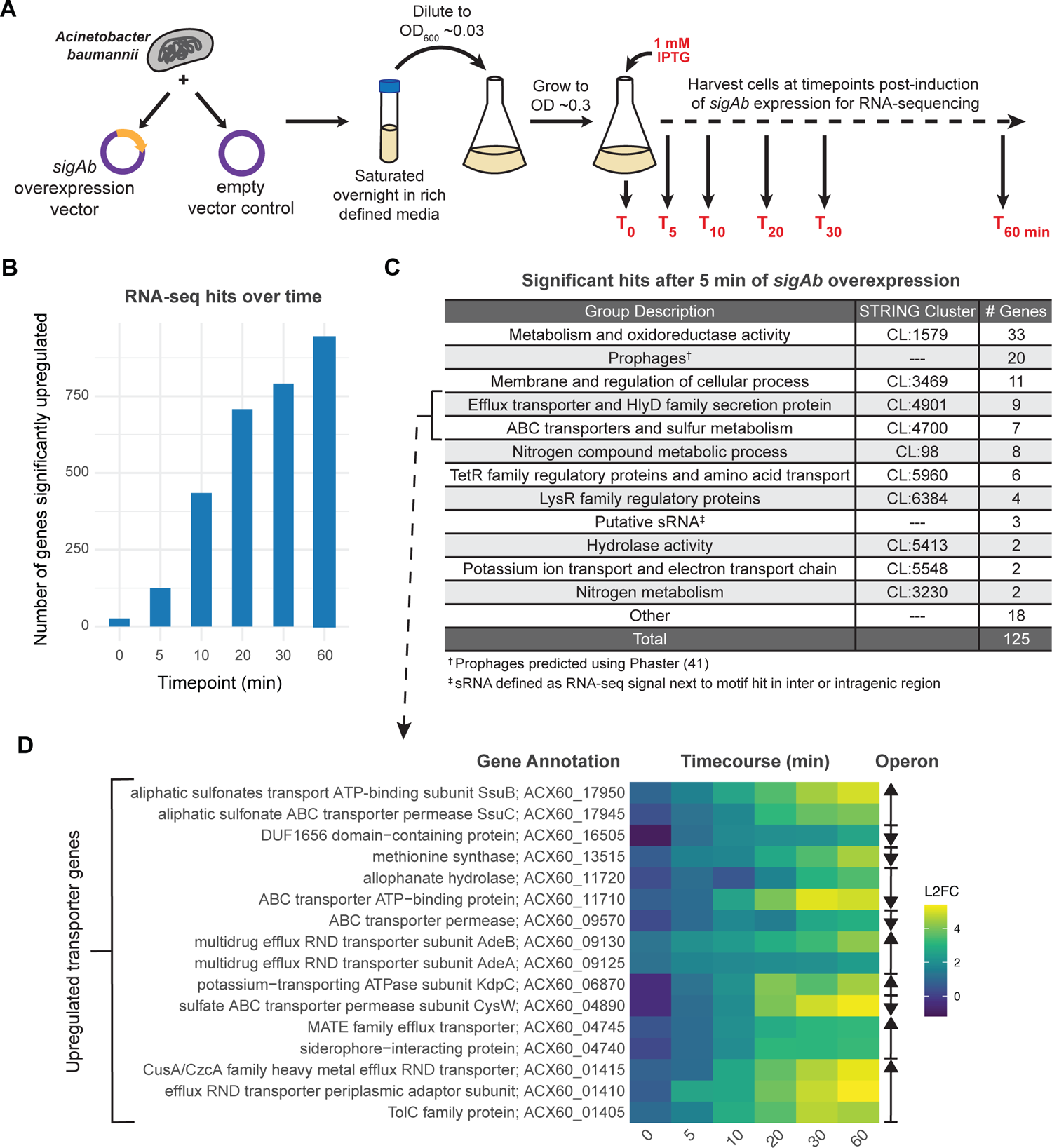
SigAb indirectly affects global transcription. **A** RNA-sequencing experimental overview. RNA was harvested and sequenced from *A. baumannii* strains harboring an inducible *sigAb* overexpression vector or empty vector control as a time course (0 to 60 minutes post-induction). **B** Number of significantly upregulated genes (Log_2_FC > 1) after induction of SigAb overexpression compared to empty vector control. RNA-seq was performed in duplicate and a false discovery rate (FDR) cutoff of 0.05 was used. **C** Table of gene set enrichments for genes significantly upregulated after 5 min of induction. **D** Heatmap of efflux pumps and resistance genes significantly upregulated (Log_2_FC > 1, FDR < 0.05, T=5 min) in RNA-seq time course experiment. Genes displayed are members of the efflux and transporter-related STRING clusters CL:4901 and CL:4700. Operons are denoted to the right, with arrows indicating direction of transcription.

These groups included the RND multidrug efflux transporter AdeA/AdeB and the CusA family heavy metal efflux RND transporter. The only gene in this group with an upstream SigAb promoter motif is the sulfate transporter CysW, suggesting indirect regulation for most of these RND efflux transporters.

In fact, among the 17 SigAb promoter motifs validated by our reporter assay, only 6 were upregulated upon SigAb OE: *sigAb* and two downstream genes that we predict form an operon, *relA*, *cysT* and downstream gene *cysW* that form an operon, and three putative sRNAs including *sabS* (Fig. S8b). We speculate that SigAb promoters that show activity in our reporter assay but lack RNA-seq signal may produce untranslated RNAs in their native context that are subject to termination and rapid degradation (42).

We reasoned that indirect effects of SigAb overexpression could be attributable to increased levels of the global regulator, RelA. To test this hypothesis, we overexpressed RelA from a strong, IPTG-inducible promoter on a multi-copy plasmid and performed RNA-seq after 10 min of induction. Although RelA OE caused significant upregulation of 62 genes, most of the genes did not overlap with those increased upon SigAb OE (Fig. S9). Therefore, the large, indirect effect of SigAb on global transcription is not through regulation of *relA*, at least under the conditions tested.

### SigAb mitigates and responds to copper stress

Upregulation of predicted heavy metal and copper transporters upon SigAb OE (Fig. 5a) suggested that SigAb could be involved in resistance to metal toxicity. Indeed, disruption of *sigAb* was found to sensitize cells to excess copper and zinc in a transposon sequencing (Tn-seq) screen of *A. baumannii* ATCC 17978 (43). To validate and extend these findings, we generated a CRISPR interference (CRISPRi) knockdown strain of *sigAb* in *A. baumannii* ATCC 19606 and phenotyped it in various transition metals. We found that *sigAb* knockdown sensitized cells to copper and nickel toxicity (Fig. 5b and S10a). We first tested liquid medium growth of the *sigAb* knockdown strain in elevated copper and nickel, finding that *sigAb* showed reduced growth relative to the non-targeting control in both conditions. To expand our phenotyping to additional conditions (e.g., manganese, cobalt), we tested growth of the *sigAb* knockdown in Biolog Phenotype Microarray (PM) plates (Fig. 5c and S10b). The PM plates recapitulated our copper and nickel results but showed no additional phenotypes. This suggests that the SigAb-dependent metal resistance is restricted to certain transition metals.

**Figure 5.**
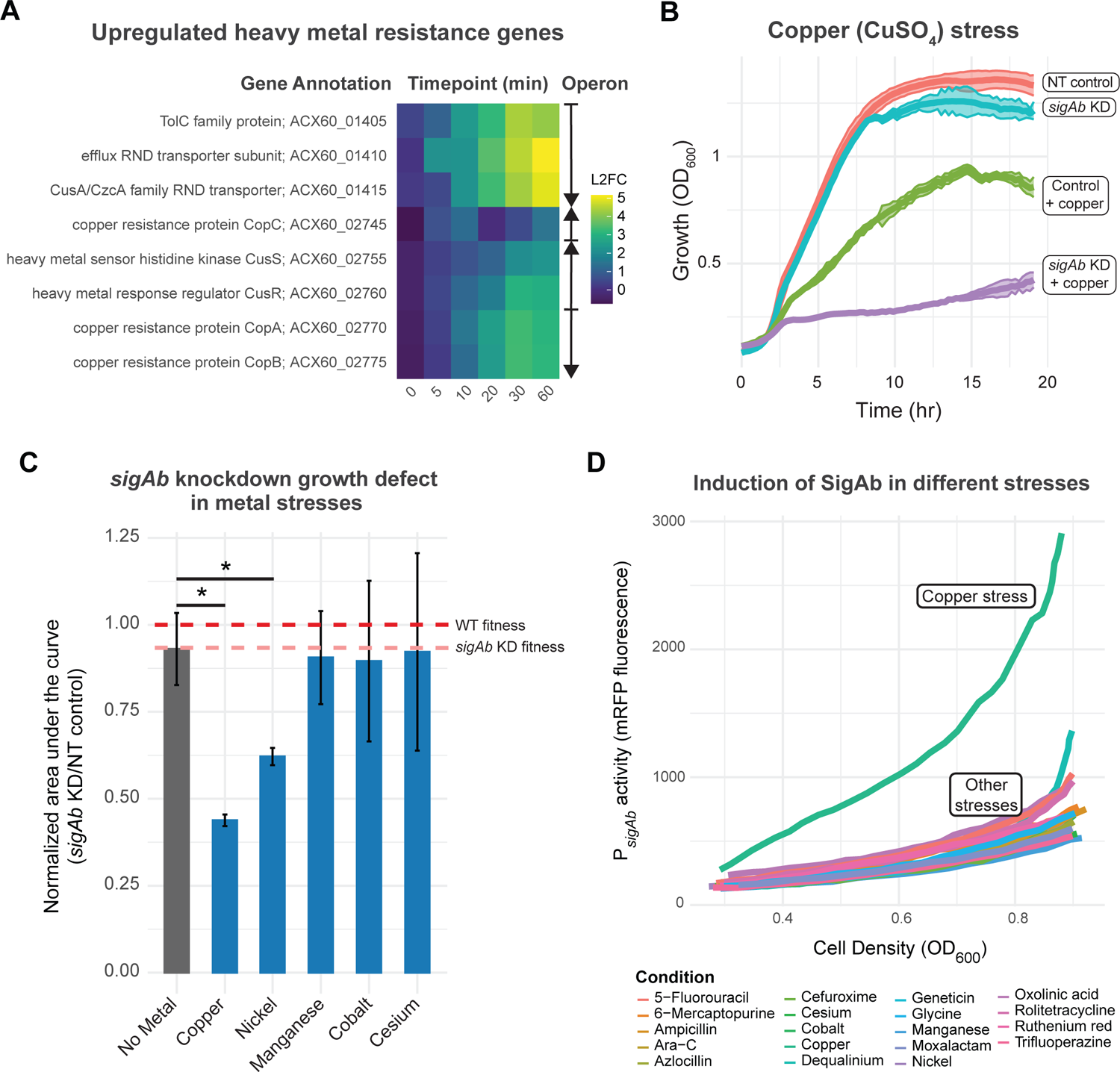
SigAb mitigates and responds to copper stress. **A** Heatmap of heavy metal resistance genes significantly upregulated (Log_2_FC > 1, FDR < 0.05, T=10 min) in RNA-seq time course experiment of *sigAb* overexpression strain compared to empty vector control. Operons are denoted to the right, with arrows indicating direction of transcription. **B** Growth curves plotted as OD_600_ over time (hr) of CRISPRi *sigAb* knockdown (KD) strain and non-targeting (NT) control in rich defined medium with 250 μg/mL CuSO_4_ stress (n=3). Data are represented as mean ± s.d. for NT control with no stress (red), *sigAb* KD with no stress (blue), NT control with copper stress (green), and *sigAb* KD with copper stress (purple). **C** *sigAb* KD growth defects in metal stresses graphed as area under the curve (AUC) normalized to NT control (n=2-9). Data are represented as the mean ± s.d. and significance was calculated with a two-tailed Student’s *t*-test (p<0.05). Bars without asterisks are not significantly different from the control **D** SigAb induction curves plotted as P*_sigAb_* activity (mRFP fluorescence) vs. cell density (OD_600_) for metal and antibiotic stress conditions using Biolog Phenotype Microarray PM13.

As small RNAs are often global regulators, we reasoned that *sabS* could be playing a role in this SigAb-dependent metal resistance. We constructed a *sabS* CRISPRi knockdown strain in *A. baumannii* ATCC 19606 and phenotyped it using Biolog PM plates. Indeed, we found that the *sabS* KD strain is sensitive to both copper and nickel stress, similar to the *sigAb* KD strain (Fig. S10c). Because SigAb activates *sabS* expression (Fig. S6b), we suggest that *sabS* is either directly or indirectly modulating SigAb-dependent metal resistance effects.

Given the role of SigAb in metal resistance, we considered that elevated metal levels could directly or indirectly stimulate SigAb activity. To test this hypothesis, we measured the activity of our P*_sigAb_* mRFP reporter under metal stress conditions using Biolog PM plates. We found that copper stress increased SigAb activity by ∼2.5-3.5-fold relative to a constitutive promoter, depending on cell density (Fig. S11a). By contrast, other metals and conditions failed to stimulate P*_sigAb_* activity above basal levels (Fig. 5d and S11b). Taken together, we conclude that SigAb activity is required for copper resistance and that SigAb responds to copper stress.

### Members of the *sigAb* operon, *aabA* and *aabB*, have anti-σ activity

We found that *sigAb* forms an operon with two uncharacterized downstream genes we call *aabA* (anti-SigAb A, ACX60_04560 in ATCC 17978) and *aabB* (anti-SigAb B, ACX60_04555 in ATCC 17978), as the coding sequences for *sigAb-aabA* and *aabA-aabB* overlap and *sigAb-aabA-aabB* are co-regulated by SigAb in our RNA-seq data (Fig. S8b). Anti-σ factors are often co-transcribed in operons with their cognate ECF σs, forming a negative regulatory loop that prevents toxicity from runaway positive autoregulation by the ECF (12). For instance, *rpoE* exists in an operon with genes that encode the anti-RpoE factor, RseA, and the RseA stabilizing protein, RseB. RseA binds to RpoE and anchors it to the membrane while RseB binds to RseA and stabilizes it against degradation by membrane proteases under non-inducing conditions (17, 44). Consistent with an RpoE-type regulation scheme, the predicted localizations of SigAb, AabA, and AabB are cytoplasmic, transmembrane, and periplasmic, respectively (Fig. S12a and S12b). We used Alphafold multimer to predict possible interactions between SigAb, AabA, and AabB, finding that AabA could bind to both SigAb and AabB *in silico* (Fig. 6a). AabB was predicted to fold around the periplasmic end of AabA, adopting a tighter alpha-helical structure than when modeled alone (Fig. S12c). AabA modeling showed a distinct interaction with SigAb compared to the interaction of RpoE and RseA, which was expected given that AabA is a much smaller protein than RseA (107 aa versus 217 aa, respectively) (Fig. 6b). Interestingly, the AabA-SigAb interaction is similar to that of the anti-sigma CnrY with sigma CnrH, a cobalt-nickel resistance regulator from the β-proteobacterium *Cupriavidus metallidurans* (Fig. 6b, (45)).

**Figure 6.**
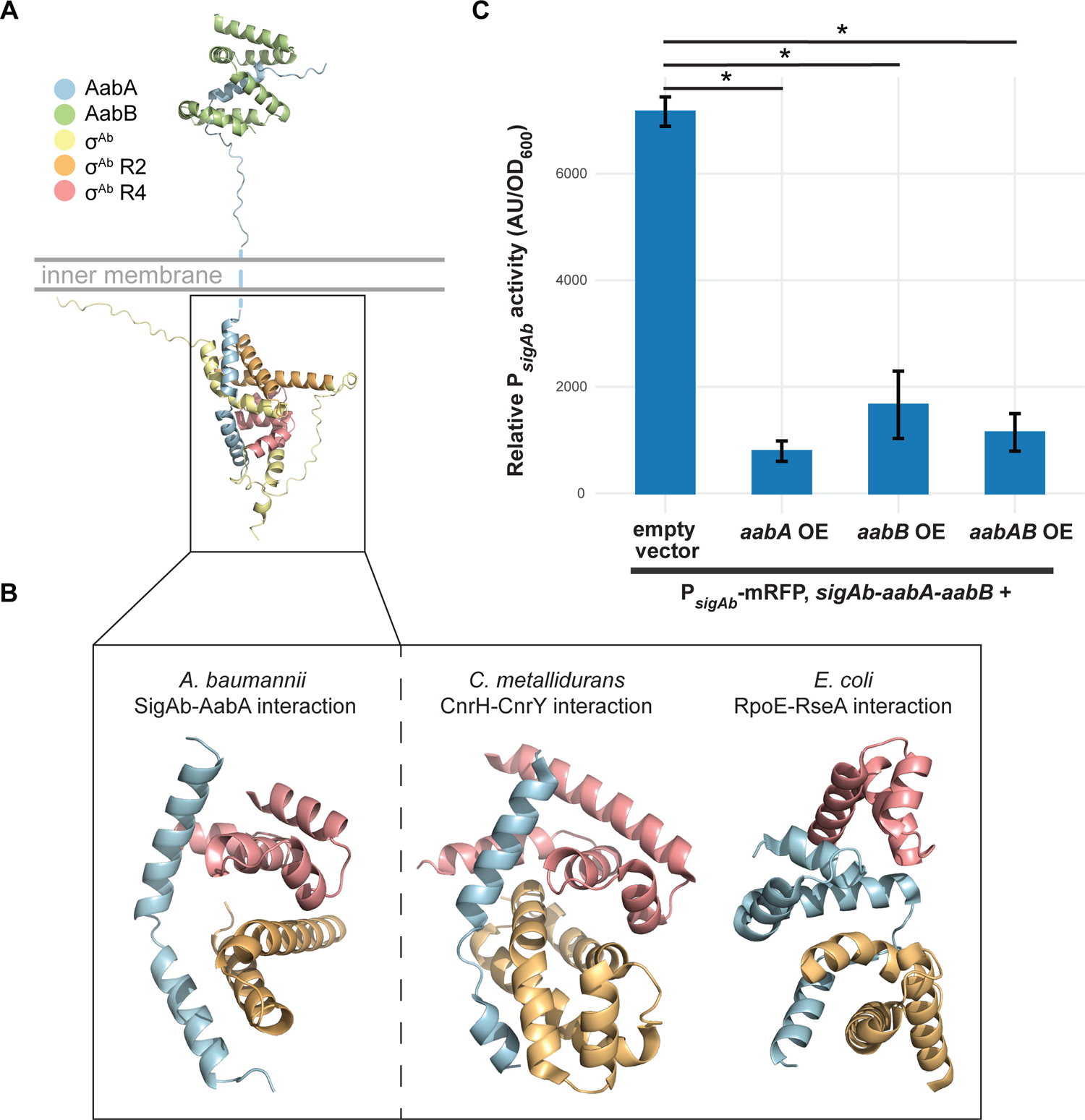
AabA and AabB have antisigma activity. **A** SigAb-AabA-AabB structural interaction model. Model predicted using AlphaFold2 run on COSMIC2 cloud platform. SigAb (σ^Ab^) region 2 (R2) and region 4 (R4) were predicted using InterProScan and AabA transmembrane domain was predicted by TMHMM 2.0. **B** Comparison of (left) *A. baumannii* σ^Ab^-AabA interaction model to (right) *C. metallidurans* CnrH-CnrY and *E. coli* σ^E^-RseA crystal structures (98, 45). AabA spans across the σ^Ab^ R2 and R4 regions, similar to CnrH-CnrY, while RseA and MucA are found in between σ^E^ R2 and R4. **C** mRFP reporter assay for P*_sigAb_* activity in *A. baumannii* strains harboring overexpression vectors with *aabA*, *aabB*, both *aabA* and *aabB*, or an empty vector control. Promoter activity is calculated as absorbance units (AU) normalized to OD_600_ (n=3). Data are represented as the mean ± s.d. and significance was calculated with a two-tailed Student’s *t*-test (p<0.05).

To test for anti-SigAb activity, we overexpressed AabA and AabB, either individually or in combination, in an *A. baumannii* strain containing our P*_sigAb_*mRFP reporter; this strain also contained the wild-type *sigAb-aabA-aabB* operon at its native locus (Fig. 6c). We found that all OE strains showed significant reduction of SigAb activity, and that activity was reduced to a similar level across strains. Although it was clear from this result that OE of *aabA*, *aabB*, or both caused anti-SigAb activity, the presence of native copies of both genes complicated interpretation of their biological roles. To eliminate interference in our assay by native *A. baumannii* proteins, we heterologously expressed SigAb, AabA, AabB and combinations thereof in an *E. coli* strain containing our P*_sigAb_* mRFP reporter (Fig. S12d). We found that co-expression of AabA and AabB significantly reduced P*_sigAb_*activity, consistent with our RpoE-like model of anti-σ function. Expressing the cytoplasmic domain of AabA alone resulted in potent inhibition of SigAb activity, demonstrating that periplasmic localization is not required for AabA activity.

Unexpectedly, we observed a significant reduction in SigAb activity when AabB was expressed alone, suggesting additional complexity to AabA-AabB anti-σ function beyond the *rpoE* paradigm. We conclude that AabA and AabB have anti-SigAb functions, although their precise mechanisms remain unknown.

### Targeted Tn-seq reveals that the *sigAb* operon is required for fitness in rich medium

The physiological importance of genes in the *sigAb* operon under standard growth conditions is largely unknown. Tn-seq studies of *A. baumannii* ATCC 17978 and AB5075 have described *sigAb* and *aabB* as non-essential and *aabA* as essential (29, 46), and a Tn disruption of *sigAb* was recovered in the ordered AB5075 mutant library. However, genome-scale Tn-seq studies can have limited resolution at the single gene level—especially for short genes (47).

Moreover, arrayed mutant libraries can accumulate secondary mutations during passaging that alter phenotypes (48). To better understand the physiological roles of *sigAb* operon genes and to establish a higher-resolution approach to gene phenotyping, we employed a CRISPR-associated transposon (CAST) system to programmatically disrupt target genes we call, “CRISPRt”. Our previously developed CRISPRt system (49) uses vectors that do not replicate in recipient bacteria to transiently express *cas* and *tns* genes from the well characterized *Vibrio cholerae* CAST (*Vc*CAST) and guide RNAs (gRNAs) with spacers that match target genes. The Cas-Tns-gRNA complex binds to target DNA complementary to the spacer (protospacer), then inserts DNA between Tn*6677* ends ∼49 bp downstream of the protospacer (Fig. 7a). Because the delivery vectors are non-replicative, insertion of Tn*6677* can be directly selected for using an antibiotic marker, similar to standard Tn-seq libraries using Tn*5* or *mariner*. We previously demonstrated that CRISPRt could inactivate reporter genes in *E. coli* K-12 (49). To establish that CRISPRt could be used for targeted Tn-seq in *A. baumannii*, we made pooled libraries of gRNA spacers targeting known non-essential (*rpoN*) and essential (*rpoD*) genes (Fig. 7b). As expected, we were able to disrupt the non-essential *rpoN* gene with Tn*6677* insertions across the entirety of the coding sequence. By contrast, targeting of the essential *rpoD* gene only allowed for insertions flanking the coding sequence, validating CRISPRt targeted Tn-seq as a high-resolution approach to determine gene essentiality in *A. baumannii*.

**Figure 7.**
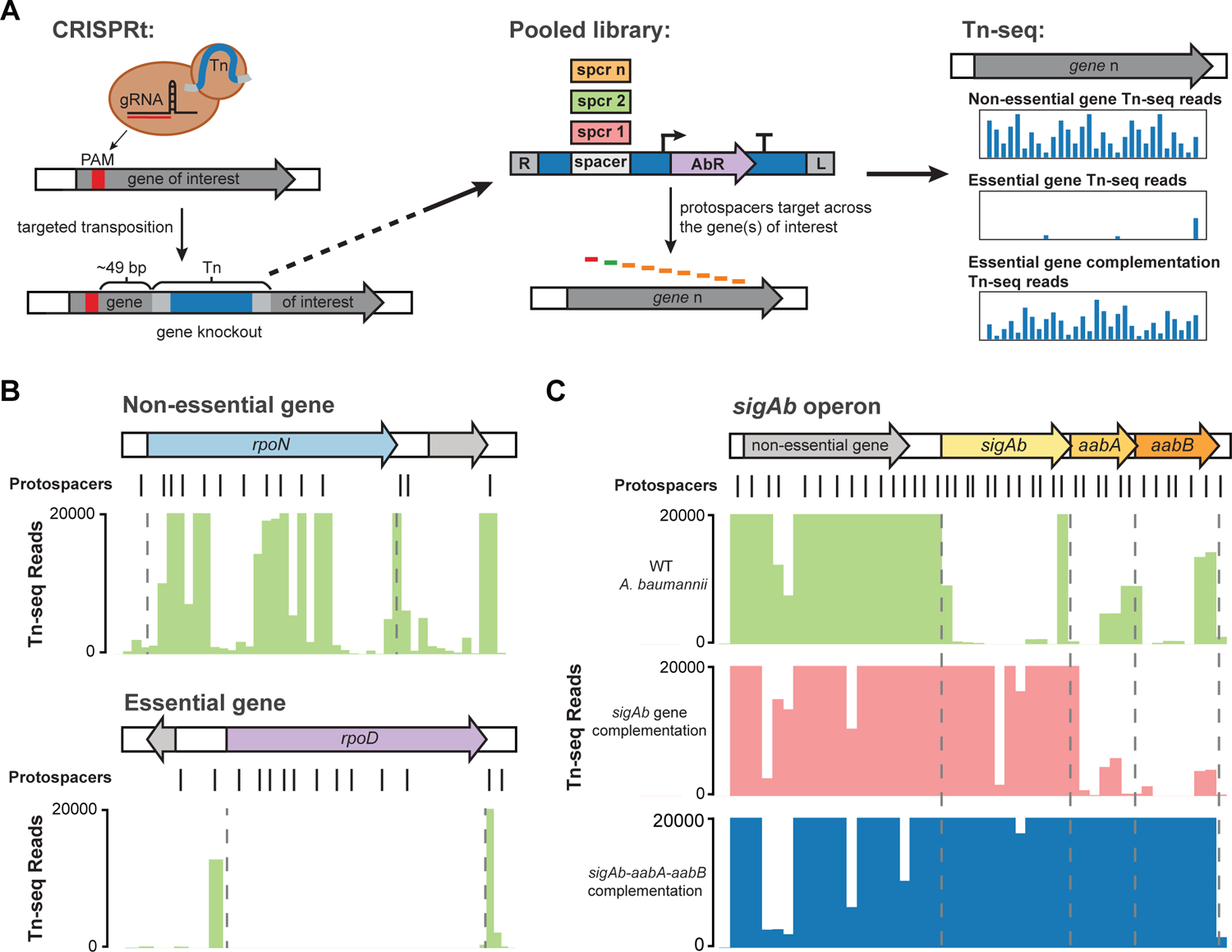
Targeted Tn-seq reveals that the *sigAb* operon is required for fitness in rich medium. **A** Schematic of CRISPR-guided targeted transposition (CRISPRt) system for gene knockouts. A high-density CRISPRt library targeting across several genes was constructed and used for gene essentiality testing. **B** CRISPRt insertions (Normalized Tn-seq reads) within the *rpoN* (non-essential) or *rpoD* (essential) genes are shown as green bars on a linear scale. Read counts are cut off at 20,000 reads due to over-representation of some insertion sites (> 250,000 reads). **C** CRISPRt insertions within the *sigAb* operon in either WT *A. baumannii* (green), a strain harboring *sigAb* gene duplication in *att*_Tn*7*_ site (pink), or a strain harboring *sigAb* operon duplication in *att*_Tn*7*_ site (blue). Normalized Tn-seq reads are shown on a linear scale with read counts cut off at 20,000 reads due to over-representation of some insertion sites (> 600,000 reads).

We next used CRISPRt to investigate essentiality of the *sigAb* operon. We tiled all three genes in the *sigAb* operon with targeting spacers as well as a predicted non-essential upstream gene (ACX60_04570) as a control. We found that Tn*6677* could be inserted across the operon, but we obtained far fewer reads from Tn insertions in *sigAb*, *aabA*, and *aabB* relative to the control gene (Fig. 7c and Fig. S13). Reduced reads from *sigAb* operon insertions could be attributed to reduced fitness of Tn insertion mutants or lower CRISPRt guide efficacy for spacers targeting the *sigAb* operon. To disambiguate these two possibilities, we performed CRISPRt Tn-seq assays on strains with a second copy of either *sigAb* or *sigAb-aabA-aabB* transcribed from their native promoter and integrated in single copy at the *att*_Tn*7*_ locus (Fig. 7c and Fig. S13).

*Trans* complementation of *sigAb* resulted in a substantial increase in *sigAb* Tn insertion reads (>30-fold) but reads for Tn insertions in *aabA* or *aabB* remained low. *Trans* complementation of the entire *sigAb* operon resulted in increased Tn-seq reads across all three genes (>30-fold), ruling out low efficiency of CRISPRt gRNAs as an alternative hypothesis. We conclude that genes in the *sigAb* operon, while not strictly essential, are required for fitness even in the absence of metal stress.

## Discussion

Elucidating the regulatory pathways by which bacterial pathogens mitigate stress may reveal new weaknesses that can be exploited by future treatments. This work substantially advances our understanding of gene regulation in the Gram-negative pathogen, *A. baumannii*, by defining the physiological roles of the ECF σ factor, SigAb. We find that SigAb is *Acinetobacter*-specific (Fig. 1) and determine that the promoter sequences it recognizes are distinct from other, well characterized σ factors (Fig. 2). By identifying SigAb direct binding sites (Fig. 3) and changes in the transcriptome during SigAb overexpression (Fig. 4), we establish a small direct and a large indirect regulon. We show that SigAb function is required for resistance to excess copper and that SigAb activity is stimulated by copper (Fig. 5), suggesting a coherent regulatory scheme for mitigating copper stress. Finally, we demonstrate that downstream genes in the *sigAb* operon have anti-SigAb activity (Fig. 6) and that disruption of any member of the *sigAb* operon leads to reduced fitness in rich medium (Fig. 7). Our work supports a growing body of literature that distinguishes regulatory strategies used by *A. baumannii* from well-studied Gram-negatives, such as *E. coli* and *P. aeruginosa*. Such distinctions may be relevant in the search for *A. baumannii*-specific treatments.

Our results further highlight fundamental differences in gene regulation strategies employed by *A. baumannii* versus related, Gram-negative pathogens. Despite previous annotations based on extrapolations from *E. coli* and *P. aeruginosa* (25), we definitively show that the only ECF σ factor present in many strains of *A. baumannii*, SigAb, is not RpoE. Both the SigAb promoter and regulon are distinct from RpoE, and RpoE-dependent promoters from *E. coli* are inactive in *A. baumannii*. Taken together with the fact that *A. baumannii* lacks other conserved σ factors, including RpoS, our results and the work of others (26, 27, 43) point to a global rewiring of gene regulatory networks that occurred sometime between the last common ancestor of *P. aeruginosa* and *A. baumannii*. As RpoE has a large, conserved direct regulon in many γ-proteobacteria (18, 22, 50, 51), and conserved genes that are part of the RpoE regulon in *E. coli* and *P. aeruginosa* are not controlled by an ECF σ in *A. baumannii*, other transcription factors must control the outer membrane stress response in *A. baumannii*. The BfmRS two-component system is one such player (27, 28), but other systems are likely involved that have not been described to date and warrant future studies.

The direct regulon of SigAb seems to contain only a handful of genes, which is consistent with many other ECF σ factors (but not RpoE) (51). By the conservative criteria applied here—namely, that direct targets must have promoter motifs, ChIP binding sites, and be upregulated by SigAb overexpression—we find only three direct targets. Autoregulation of the *sigAb-aabA-aabB* operon at the transcriptional level is a hallmark of ECF σ factors (8, 51), and the presence of a negative feedback loop consisting of an anti-σ factor is also commonplace (12, 52). The smaller size of AabA compared to RseA and substantial anti-σ activity of the AabA cytoplasmic fragment raise questions about AabA proteolysis and release of SigAb, possibly due to a copper stimulus. Direct control of the stringent response factor, *relA* (37), by SigAb has unknown functional consequences. RNA-seq of RelA or SigAb overexpressing strains showed little overlap, seemingly ruling out increased RelA/(p)ppGpp as a cause of the indirect regulon. However, there may be other conditions in which the regulatory relationship between SigAb and RelA plays a functional role, such as during metal stress in the host environment. Another host-associated pathogen, *Mycobacterium tuberculosis*, also uses an ECF σ factor to control expression of *relA* (53, 54), suggesting a functional convergence between these two unrelated pathogens. Finally, we show that SigAb directly regulates the putative small RNA, *sabS*. This sRNA was previously detected in a transcriptome study (39), but its dependence on SigAb was unknown until our work. A recent preprint displaying GRIL-seq data in *A. baumannii* suggests that the downregulated genes in our *sigAb* overexpression RNA-seq dataset are direct targets of *sabS*, however these genes are mostly unannotated, hypothetical proteins (55). sRNAs are often core members of ECF σ regulons (56, 57). By example, the essential function of RpoE in *E. coli* is to transcribe three sRNAs that downregulate key envelope target genes (20, 58). Of the direct SigAb targets, SabS is most likely to be responsible for the changes seen in the indirect regulon, given that *sabS* and *sigAb* knockdowns show similar phenotypes. However, how SabS can increase the expression of hundreds of genes is unknown and may be due to additional, indirect regulation. The indirect regulon of SigAb is critical to its physiological roles. Our phenotyping experiments point to SigAb, and by extension, SabS, as important mediators of the metal stress response, and we find that SigAb indirectly regulates several heavy metal RNA efflux pumps.

Metal stress and acquisition is key to pathogenesis by *A. baumannii* and, thus, SigAb may be important under those conditions. The levels of transition metals, such as copper and manganese, are tightly regulated during growth in the host environment, as bacteria require metals for viability but metals are toxic in excess (59, 60). Copper may be relevant for other clinical settings as it is often used as a bactericide due to its high toxicity to bacteria (61). *A. baumannii* is known to evade copper toxicity via several gene clusters including the resistance genes, *copA*/*copB*, (62) as well as the histidine kinase-response regulator pair, *cusRS* (63); these genes are also part of the SigAb indirect regulon. Interestingly, SigAb is distinct from other copper responsive ECF σ factors that use an anti-σ-independent mechanism for signal transduction (ColE-like ECFs) or through activation of carotenoid genes in response to copper stress (CarQ-like ECFs) (64). Taken together, the role of SigAb in mitigating metal stress underscore its importance to *A. baumannii* physiology and potentially pathogenesis.

In this study, we used a targeted version of Tn-seq called “CRISPRt” to characterize gene fitness and essentiality. CRISPRt has advantages for probing the essentiality of subsets of genes that may be broadly applicable. For instance, Tn-seq using pseudo-random insertions (e.g., Tn*5*) often fails to achieve high-density coverage of specific loci unless very large insertion libraries are constructed. The need to construct such libraries typically precludes the use of genetic complementation in the context of Tn-seq; however, CRISPRt targeting of specific loci makes Tn-seq complementation possible with small libraries. Targeted, CRISPRt follow-ups of specific loci could provide a way to validate essential gene calls from pseudo-random Tn-seq at scale, which would be especially valuable for non-model bacteria. Importantly, our CRISPRt analysis had sufficient resolution to determine that members of the *sigAb* operon had reduced fitness, in contrast to other studies that made binary essential/non-essential calls (29, 46). As with pseudo-random transposition, polar effects onto downstream genes from changes in transcription are a concern, but the possibility of complementation mitigates that issue, and the trade-off may be worthwhile for large-scale studies validating dozens of genes at a time. Demonstrating CAST-based, targeted Tn insertion in *A. baumannii* adds another genetic tool for gene phenotyping to this urgent threat pathogen.

## Materials and Methods

### Strains and growth conditions

Strains are listed in Table S1. *Escherichia coli* K-12 and *Acinetobacter baumannii* (strains ATCC 17978-UN or ATCC 19606) were grown in Lennox lysogeny broth (LB) at 37°C shaking in a flask at 250 rpm, in a culture tube on a rollerdrum at max speed, in a 96-well plate shaking at 900 rpm, or in a plate reader shaking (Tecan Infinite Mplex, Infinite Nano+, or Sunrise). Culture medium was solidified with 1.5% agar for growth on plates. Where noted, *A. baumannii* strains were grown in EZ Rich Defined Medium (Teknova M2105), following manufacturer’s recipe except supplemented with 40 mM succinate instead of glucose (AbRDM). Antibiotics were added when necessary: 100 μg/mL ampicillin (amp), 30 μg/mL kanamycin (kan), 50 μg/mL apramycin (apr), 50 µg/mL spectinomycin (spec) for *E. coli* and 150 μg/mL carbenicillin (carb), 60 μg/mL kanamycin (kan), 100 μg/mL apramycin (apr), 150 μg/mL gentamycin (gent) for *A. baumannii*. Diaminopimelic acid (DAP) was added at 300 μM to support growth of *E. coli* dap-donor strains. IPTG (isopropyl b-D-1-thiogalactopyranoside) was added at 0 to 1 mM as indicated in the figures or figure legends. Strains were preserved in 15% glycerol at −80°C. Plasmids were propagated in *E. coli* strain BW25141 *att*_Tn*7*_::*acrIIA4* (sJMP3053) for DNA extraction and analysis or in *E. coli* strain WM6026 *att*_Tn*7*_::*acrIIA4* (sJMP3257) for conjugation.

### General molecular biology techniques

A complete list of plasmids and oligonucleotides are listed in Tables S2 and S3. Oligonucleotides were synthesized by Integrated DNA Technologies (Coralville, IA). Genomic DNA was purified using GeneJet Genomic DNA kit (Thermo K0503). Plasmid DNA was purified using the GeneJet Plasmid Miniprep kit (Thermo K0503) or the Purelink HiPure Plasmid Midiprep kit (Invitrogen K210005). PCR was performed according to manufacturer directions using Q5 or OneTaq DNA Polymerases (NEB, Ipswitch, MA). DNA was digested with restriction enzymes from New England Biolabs (NEB). PCR products were purified with DNA Spin and Concentrate kit (Zymo Research, Irvine, CA, D4013 or NEB Monarch, T1030) following manufacturer instructions or gel-purified from kit (Zymo Research). Plasmids were assembled using NEBuilder HiFi DNA assembly kit (NEB). DNA was quantified on a Nanodrop Lite or Qubit HS DNA or RNA kit (Thermo). Plasmids were transformed into electrocompetent *E. coli* cells using a 0.1 cm cuvette (Fisher FB101) and a BioRad Gene Pulser Xcell (25 μF, 200 ohm, 1800 V). Plasmids and recombinant strains were sequenced via Sanger sequencing by Functional Biosciences or Oxford Nanopore sequencing by Plasmidsaurus. Next-generation sequencing was performed by the UW-Madison Biotechnology Center Next Generation Sequencing Core using an Illumina NovaSeq 6000 or Azenta using an Illumina MiSeq.

### SigAb structural modeling

The SigAb protein sequence was structurally modeled using Phyre2 “Normal” setting to identify proteins with similar structures (See Fig. S1). To model based off of *E. coli* RpoE specifically, Phyre2 was used with “one-to-one threading” (33). Structural model was traced onto RpoE holoenzyme crystal structure (34) in PyMOL.

### Evolutionary analysis of SigAb

SigAb targeted ortholog search and phylogentic analyses were constructed similar to that previously described (65). Briefly, 2822 γ-proteobacteria genome assemblies were retrieved from the RefSeq database (release 213, downloaded on September 2022), selecting reference genomes and isolates of interest, including 196 isolates from the *Acinetobacter* genus. To perform phylogentic profiling, protein reference sequences of RNA polymerase sigma factors were selected from *Acinetobacter baumannii* ATCC 19606 (WP_000362312.1) and *Escherichia albertii* Sample 167 (WP_001295364.1). Phylogenetic profiles were computed using fDOG 0.1.23 (https://github.com/BIONF/fDOG) with the following parameters: compilation of 35 core orthologs selected with a taxonomic distance minimum of genus and maximum of class.nHomologous sample protein sequences were aligned with MAFFT (66) and reconstructed into a phylogeny using FastTree (67). Aligned sequences were classified into clades in a phylogenetic-aware manner using Fastbaps (68).

### 5’ RACE

The 5’ end of the sigAb transcript was identified using 5’ RACE following manufacturer’s protocol with template switching RT enzyme mix (NEB; M0466). Briefly, cDNA from *A. baumannii* ATCC 17978 RNA was made using RT oligo oJP2139 and template-switching oligo oJMP2131. 5’ region of sigAb transcript was amplified with oJMP2130/oJMP2138 using NEBNext Ultra II Q5 master mix and touchdown PCR, spin purified with DNA clean and concentrate kit (NEB), and sequenced with Plasmidsaurus.

### Promoter activity fluorescent assays

Putative P*_sigAb_* motifs were cloned into Tn*7*-based mRFP reporter vectors using annealed oligos and ligation of 54 bp promoter motifs into BsaI-cut vector (pJMP3570). These promoter reporters were integrated onto the chromosome in the *att*_Tn7_ site in sJMP3075 (*E. coli* MG1655 WT), sJMP3348 (*A. baumannii* ATCC 17978 WT), and sJMP3329 (*A. baumannii* ATCC 19606 WT) by tri-parental Tn*7*-mediated conjugation with the transposase vector-containing donor strain, sJMP3261. Strains were grown in AbRDM in a 96-well plate overnight to saturation and red fluorescence and OD_600_ were measured in a Tecan Infinite Mplex or Nano+ plate reader.

### Promoter Mutagenesis Screen

A P*_sigAb_* mutation library containing 54 bp promoter region with individual point mutations in putative −10 and −35 motifs, and chunk mutations in putative UP element and spacer region T-tract, was constructed as follows. An oPool (oJMP2304) containing 66 oligos with the mutated promoter region was amplified with oJMP463/oJMP464 using low-cycle Q5 PCR with the following conditions: 98°C 30s; 98°C 15s, 56°C 15s 72°C 15s, 16 cycles; 72°C 5 min. PCR product was spin-purified, quantified with nanodrop, digested with PacI/SpeI, ligated into PacI/SpeI-digested pJMP3539, and transformed into the mating strain to make sJMP3544 containing the P*_sigAb_* mutation library. The PsigAb library was integrated into the attTn7 site in *A. baumannii* ATCC 19606 by quad-parental mating with sJMP3329 (WT *A. baumannii*) + sJMP4061 (helper plasmid) + sJMP3261 (Tn*7* transposase) + sJMP3544 (P*_sigAb_* mutation library) and selection with apr. Isolated colonies were picked into 3 96-well plates, grown up in LB + apr overnight to saturation, and stored as sJMP3565, sJMP3566, and sJMP3567.

To determine the identity of each promoter mutation in each well, barcoded colony PCR followed by sequencing was performed as follows. Cells in each 96-well plate were diluted 1:100 and 2 µL were added to OneTaq PCR mix with oJMP2292/oJMP1678-1773 containing 6 nt defined barcodes. Barcoded PCR products from each plate were pooled together, spin purified, and sequenced. The identity of each mutation in each well was decoded using the barcodes as a key. Once the identity of each well was found, the P*_sigAb_* promoter activities were determined by growing up the 96-well plates in AbRDM to saturation and quantifying red fluorescent protein normalized to OD_600_ in a Tecan Mplex. The median activity for each mutation was compiled into an activity logo using WebLogo sequence logo generator (69).

### Chromatin Immunoprecipitation-sequencing (ChIP-seq)

An expression vector harboring N-terminally tagged SigAb was constructed by amplifying HaloTag (HT) gene from sJMP3331 gDNA using oJMP2295/oJMP2296 with a gly-ser-gly-ser flexible linker and no translation stop codon, amplifying *sigAb* from sJMP3348 gDNA using oJMP2294/oJMP1905, and HiFi assembling into NcoI/BamHI-digested pJMP3653 expression vector containing a strong, IPTG-inducible promoter (70) to make pJMP3571. pJMP3571 was electroporated into the mating strain (sJMP3257) to make sJMP3575. To make an *A. baumannii* ATCC 17978 strain containing the HT-SigAb expression vector (kanR), sJMP3575 was mated with sJMP3348 to make sJMP3584. ChIP-seq on sJMP3584 was performed in triplicate as described previously (40, 71).

Briefly, cells were grown in 100 mL AbRDM + 70µM IPTG + kanamycin to maintain the plasmid until reaching mid-log (∼OD 0.3). Cultures were crosslinked with formaldehyde, quenched with glycine, and 50 mL cell pellets were harvested. Pellets were sonicated in a Covaris Misonix sonicator for 16 min (20% duty factor, 75 PIP, 200 cycles per burst, 6°C) to achieve ∼100-500 bp fragments and immunoprecipitation of HaloTagged-SigAb protein was performed according to Promega HaloChIP protocol. Before IP, 1/10 of each sample was saved as input control. Input control and HT-enriched samples were prepared for Illumina sequencing using NEBNext Ultra II Library Prep Kit for Illumina (NEB; E7645S) following manufacturer’s protocol and sequenced on Illumina NovaSeq 6000 2×150 with the UW-Madison Biotechnology Center at ∼10 million reads per sample.

ChIP-seq paired end FASTQ files were filtered to remove low quality bases using Trimmomatic (72) (v0.3) (Sliding window of 3:30, Minimum length of 36 bp, leading and trailing both a value of 3) and aligned to the *Acinetobacter baumannii* ATCC 17978 genome (GCA_001077675.1) using Bowtie2 (v2.2.2) (73) and default parameters. Samtools (v1.2) (74) and Picard Tools (v1.98) (75) were used to convert the SAM file to a sorted BAM file. Deeptools (v3.5.1) (76) was used to generate IP vs INPUT ratio files for visualization (binsize of 1 and readCount scaleFactorsMethod). ChIP-seq peaks were identified with the IP and INPUT BAM files using MACS3 (v3.0.0) (77) with default parameters except for using the “nomodel” option and 128 for the “extsize” and all for the “keep-dup” values.

### RNA-sequencing

RNA-sequencing (RNA-seq) was performed on *A. baumannii* ATCC 17978 strains harboring either a *sigAb* overexpression (OE) vector (sJMP3382) or an empty vector control (sJMP3380). Cells were diluted 1:100 from a saturated overnight into AbRDM + kan to maintain the plasmid and grown up shaking at 37°C. Once reaching mid-log (OD 0.3), 1 mM IPTG was added to induce expression, and cell pellets were collected for RNA purification at timepoints from 0 to 60 min after addition of IPTG. For *relA* OE experiments, *A. baumannii* ATCC 17978 strains containing *relA* OE vector (sJMP3790) or empty vector control (sJMP3719) were grown as described above, except cell pellets were collected for RNA purification at timepoints 0 and 10 min only.

Total RNA was extracted from *A. baumannii* using hot phenol organic extraction, as previously described (78). Briefly, mid-log cells were added to 1.25 mL stop solution (5% water-saturated phenol in ethanol), spun down at 11,000xg for 5 min at 4°C, and pellets were flash frozen in a dry ice:ethanol bath and stored at −80°C. Cells were lysed with lysozyme and SDS, total RNA was purified with phenol, phenol:chloroform, and chloroform extractions followed by ethanol precipitation, and residual DNA was removed with Turbo DNase I treatment (Invitrogen). Ribosomal RNA (rRNA) was depleted from the total RNA samples as previously described (79). Briefly, DNA oligos complementary to *A. baumannii* 23S, 16S, and 5S rRNA were annealed to the total RNA samples, RNase H treatment was performed to cleave the annealed rRNA, and the DNA oligos were removed with Turbo DNase I. rRNA-depleted samples were prepared for next-generation sequencing using NEBNext Ultra II Directional RNA Library Prep kit (NEB; E7765S) and NEBNext Multiplex Oligos for Illumina (NEB; E6640S). Libraries were sequenced on Illumina NovaSeq 6000 2×150 with the UW-Madison Biotechnology Center at ∼10 million reads per sample.

Sequencing reads were trimmed using Trimmomatic (72) (version 0.39) (default parameters except for Sliding window of 3:30, Minimum length of 36 bp, leading and trailing both a value of 3) and mapped to the *A. baumannii* ATCC 17978 genome (GCA_001077675.1) using bwa-mem (80) (version 0.7.17-r1188) using default parameters. Mapped reads were further processed with Picard-tools (version 2.25.10) (CleanSAM and AddOrReplaceReadGroups) (75) and samtools (74) (version 1.2) (sort and index). Paired aligned reads were mapped to genes with HTSeq (81) (version 0.6.0) with default parameters and normalized using FPKM as previously described (82, 83). The R package edgeR (84) (version 3.30.3) was used for differential gene expression analysis using Benjamini and Hochberg (85) adjusted P value (FDR) ≤ 0.05 as the significance threshold. Sequencing reads per gene were normalized using the fragments per kilobase per million mapped reads method (FPKM). Data were visualized in R using ggplot2.

### CRISPRi knockdown experiments

sgRNAs targeting *sigAb* and *sabS* were cloned into pJMP2776 using oligos oJMP1243/oJMP1244 and oJMP2632/oJMP2633, respectively, as previously described to make pJMP3353 and pJMP3854 (86). The CRISPRi system was integrated into the *att*_Tn*7*_ site in *A. baumannii* ATCC 19606 (sJMP3329) using quad-parental mating and selection on gentamycin as previously described (87). *sigAb* knockdown strain (sJMP3363), *sabS* knockdown strain (sJMP3856), and non-targeting sgRNA strain (sJMP6498) were assayed for KD-dependent phenotypes by growing to saturation overnight in AbRDM, then pre-depleting by diluting 1:100 in AbRDM + 1mM IPTG for 4 hours (mid-log), and finally diluting 1:100 again in AbRDM + 1 mM IPTG + chemical (as indicated in figure legends) and measured OD_600_ for 16 or 18 hours in a Tecan Sunrise, Infinite Mplex, or Infinite Nano+ plate reader. Biolog phenotyping experiments were performed with the same pre-depletion method, except the final 1:100 diluted mid-log cells with 1 mM IPTG but no additional chemical were added to each well. Biolog Phenotype Microarray plates PM13 and PM16 were used.

### SigAb induction phenotyping

SigAb promoter (P*_sigAb_*) mScarlet-I reporter strain in *A. baumannii* ATCC 19606 (sJMP3406) and constitutive promoter (P*_lacUV5_*) reporter strain (sJMP3402) were grown to saturation in LB. Saturated culture was diluted back 1:100 in AbRDM and grown up to mid-log. For Biolog Phenotype Microarray assays, 100 µL of cells diluted 1:100 in fresh AbRDM were added to each well of plates PM11, PM13, and PM16. For copper testing, cells were diluted 1:100 into AbRDM supplemented with 150 µg/mL CuSO_4_. 96-well plates were grown in Tecan Infinite Mplex or Nano+ plate readers and measured OD_600_ and red fluorescence.

### SigAb-AabA-AabB structural predictions

Structural interactions between σ^Ab^, AabA, and AabB were predicted using Alphafold2 multimer (88) run on the COSMIC2 cloud platform with the following parameters: Database: full_dbs, Model: multimer, Number of predictions per model: 1, Latest date (YYYY-mm-dd) to use for template search: 2023-05-30, Models to relax: none. Interactions between 1) SigAb and AabA or 2) AabA and AabB were predicted separately. Multimer models containing AabA used either a cytoplasmic (modeled with σ^Ab^) or periplasmic (modeled with AabB) fragment based on the location of a predicted transmembrane helix (residues 42-64, predicted by TMHMM 2.0 (89)). A predicted signal peptide in AabB (residues 1-21, predicted by SignalP 6.0 (90), was removed prior to modeling. σ^Ab^ sequences corresponding to σ_R2_ and σ_R4_ were predicted using InterProScan (91).

### Anti-SigAb phenotyping

*In A. baumannii*: *aabA*, *aabB*, or *aabA-aabB*, were cloned into pJMP3352 under control of the promoter P*_trc_* according to “construction/notes” in Table S2 to make plasmids pJMP3603, pJMP3604, and pJMP3549, respectively. Plasmids, including empty vector pJMP3352, were mated into P*_sigAb_* mRFP reporter strain (sJMP3602) to make strains sJMP3629, sJMP3630, sJMP3631, and sJMP3380. Strains were grown overnight to saturation in AbRDM supplemented with kan and 1 mM IPTG. OD_600_ and red fluorescent protein were measured in a Tecan Infinite Mplex plate reader.

*In E. coli*: *aabA*, *aabB*, *aabA-aabB*, or *aabA*-cytoplasmic domain were cloned into pJMP10740 under control of the promoter P*_araBAD_* according to “construction/notes” in Table S2 to make plasmids pJMP3802, pJMP3803, pJMP3804, and pJMP3805, respectively. Plasmids were co-mated into BW25113 P*_sigAb_*mRFP reporter strain (sJMP3821) with *sigAb* overexpression plasmid (pJMP3735) to make strains sJMP3822, sJMP3823, sJMP3824, and sJMP3825. Strains were grown overnight to saturation in AbRDM supplemented with kan, spec, 50 µM IPTG, and 10 mM L-arabinose. OD_600_ and red fluorescent protein were measured in a Tecan Infinite Mplex plate reader.

### Targeted Tn-seq (CRISPRt) experiments

SigAb operon complementation strains were constructed by amplifying the *sigAb* gene with oJMP2344/2345 and the *sigAb* operon with oJMP2344/2347, assembling into SpeI/AscI-digested pJMP8602 to make pJMP3607 and pJMP3609, and using conjugation with selection on apramycin to integrate into the *att*_Tn*7*_ site in *A. baumannii* ATCC 17978 (sJMP3348), resulting in sJMP3624 and sJMP3628. CRISPRt guides were designed to have a “CN” PAM with the spacer being 32 nt.

CRISPRt gRNA library targeting the *A. baumannii* genes *rpoN*, *rpoD*, *rpoH*, *mdcD*, *sigAb*, *aabA*, and *aabB* was constructed as follows. An oPool (oJMP2322) containing guides targeting the first 4 genes (15 guides per gene) was amplified using oligos oJMP463/oJMP464 using low-cycle PCR as described above. An oPool (oJMP2143) containing 50 gRNAs targeting the *sigAb* operon was amplified using oligos oJMP463/oJMP464 and oJMP465/oJMP466 using the same PCR conditions. PCR products were spin-purified, quantified with nanodrop, and pooled together to achieve approximately equal gRNA ratios. Pooled product was digested with BsaI, ligated into BsaI-digested pJMP10621, and transformed into the mating strain to make sJMP3576 containing the CRISPRt plasmid library.

CRISPRt targeted transposition experiment was performed using tri-parental mating overnight at 30°C of: CRISPRt libarary (sJMP3576), CRISPRt helper plasmid (sJMP10275), and recipient strains 1) WT *A. baumannii* (sJMP3348), 2) *sigAb* gene complementation (sJMP3624), or 3) *sigAb* operon complementation (sJMP3628) with selection on apramycin to create strains sJMP3693, sJMP3694, and sJMP3696, respectively. Colonies were scraped off of 150mm plates (∼5,000 colonies per strain) using LB and a cell scraper and gDNA was extracted. Tn-seq library was prepared for Illumina sequencing as previously described (49). Briefly, gDNA was cut with MmeI enzyme, adaptor oligos oJMP1995/oJMP1996 were annealed and ligated to the DNA, and a low-cycle PCR was performed with oligos oJMP1997/oJMP1998. Samples were sequenced on Illumina NovaSeq 6000 2×150 with the UW-Madison Biotechnology Center at ∼10 million reads per sample.

CRISPRt FASTQ files containing the transposon sequence (R1 FASTQ files) were trimmed to remove the transposon sequence using Cutadapt (v3.4) (92). Resulting reads longer than 40 nts were removed using fastp from Deeptools (v3.5.1) (76). Bowtie (v1.0.0) (93) was used to align the reads ≤ 40 nts using default parameters. Samtools (v1.2) (74) was used to convert the SAM file to a sorted BAM file and Deeptools (v3.5.1) (76) was used to generate BigWig files for visualization. Unique hits aligning to either strand were identified using Samtools (v1.2) and the standard Linux commands of awk, sort, and uniq to filter the alignment file to count aligned reads on the forward or reverse strands.

## Data Availability

Raw data will be deposited to the National Center for Biotechnology Information Sequencing Read Archive (SRA) under BioProject (### pending). All other data is available upon request.

## Acknowledgements

This work was supported by the National Institutes of Health under award numbers K22AI137122 and 1R35GM150487-01. We thank the UW-Madison Biotechnology Center Illumina Pilot Grant Program for providing funding for sequencing. We thank Lauren Palmer and members of Peters lab for helpful comments.

